# PARMIK: PArtial Read Matching with Inexpensive K-mers

**DOI:** 10.1101/2024.10.14.618242

**Authors:** Morteza Baradaran, Ryan M Layer, Kevin Skadron

## Abstract

Environmental metagenomic sampling is instrumental in preparing for future pandemics by enabling early identification of potential pathogens and timely intervention strategies. Novel pathogens are a major concern, especially for zoonotic events. However, discovering novel pathogens often requires genome assembly, which remains a significant bottleneck. A robust metagenomic sampling that is directly searchable with new infection samples would give us a real-time understanding of outbreak origins dynamics. In this study, we propose PArtial Read Matching with Inexpensive K-mers (PARMIK), which is a search tool for efficiently identifying similar sequences from a patient sample (query) to a metagenomic sample (read). For example, at 90% identity between a query and a read, PARMIK surpassed BLAST, providing up to 21% higher recall. By filtering highly frequent k-mers, we reduced PARMIK’s index size by over 50%. Moreover, PARMIK identified longer alignments faster than BLAST, peaking at 1.57×, when parallelizing across 32 cores.

## I. Introduction

Environmental metagenomic sampling is a pivotal tool in preparing for future pandemics by enabling the early identification of potential pathogens and facilitating prompt intervention strategies. When we encounter patient samples of unknown origin, it would be helpful to draw a connection to environmental data to discover the pathogen, especially for zoonotic events. As illustrated in the figure 1, this process starts with the collection of environmental samples, followed by a whole-genome sequencing step. Today’s high-throughput sequencing technologies like Illumina [1] can generate millions to billions of *reads*, short substrings of DNA molecules represented by base pairs (A, C, G, T). However, the subsequent data analysis step, involving assembling or identifying the reads through *alignment* with reference genomes, is notably slower. A robust and directly searchable environmental metagenomic database for newly collected patient samples would significantly enhance our ability to rapidly identify the origins of outbreaks, bypassing the slow assembly phase.

**Fig. 1.**
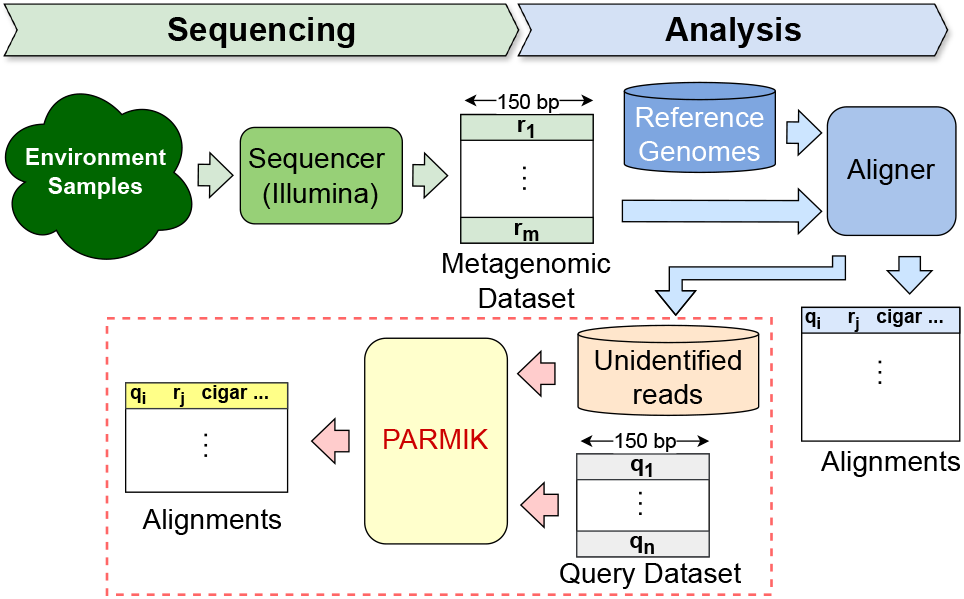
Environment Sampling, Sequencing, Alignment, and Partial Matching.

In reference genome alignment-based methods, a significant portion of reads fail to find alignment with any reference genome, thus remaining *unidentified*. This problem is primarily related to the inherent genetic diversity and novel sequences within metagenomic samples, which often lack suitable matches in existing reference databases. Hence, analysis of these unidentified metagenomic samples (red dotted rectangle in figure 1) remains an under-studied area, with no effective tool as far as we know. However, it holds significant importance in the context of pandemic preparedness, making it the main focus of our research.

One case of metagenome analysis methodologies begins by identifying specific, fixed-length substrings within the reads, often referred to as *k-mers*^*^. The underlying principle here is that if two reads are similar, they should share some k-mers. K-mer search methods, such as the well-known Kraken [13], [14], can identify well-known pathogens. However, they fall short when detecting previously undiscovered pathogens for which no reference genome exists. Although k-mer processing algorithms are diverse, they all require large storage for k-mer sets, making the reduction of storage needs and optimization of query speed critical challenges. A second class of methods based on pseudo-alignment, like Themisto [16], exhibit rapid estimation of read alignments between two sequences. While these methods can quickly indicate the presence or absence of highly similar sequences within a metagenomic dataset, they fall short of providing precise details, such as whether the matches are biologically relevant or just random, non-specific alignments, which is necessary for comprehensive epidemiological or virological analysis.

To address these shortcomings, a critical need arises for an advanced search tool. This tool should be able to identify and extract all potentially relevant reads with *“partial matches”* from hundreds of millions of unassembled and unaligned metagenomic reads quickly and with an efficient memory footprint. A partial match (figure 2) is a region between a read and a query where the boundaries of the read and query do not necessarily align, and the overlapping region can be small, including regions of exact matches and a few mismatches, substitutions and indels. Existing tools like BWA often overlook alignments below a specific size, while BLAST struggles with alignments that contain significant mismatches despite exact matches in certain segments of the alignment. When aligning Wuhan-Hu-1 to Bat Feces, BLAST, and BWA miss finding any match for 1.7% and 10.3% of queries, respectively. To our knowledge, there is currently no tool that effectively addresses these types of partial matches.

**Fig. 2.**
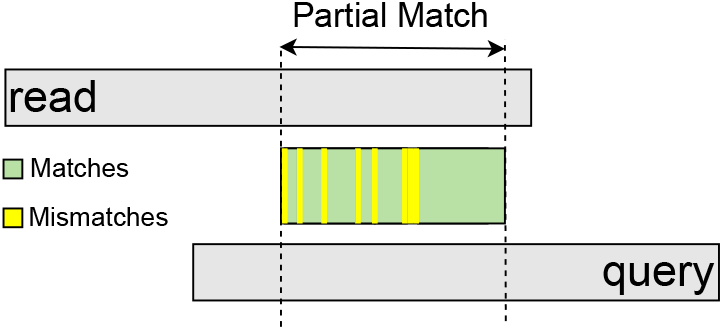
In the partial match region, the boundaries of the read and query are not necessarily aligned, and the overlap region can be small, including regions of matches and a few mismatches.

In this project, we introduce PArtial Read Matching with Inexpensive K-mers (PARMIK), a search tool designed to identify the partial match between all sequences from a patient sample (query) to a metagenomic sample (read). PARMIK employs a four-step approach: indexing, pre-filtering, alignment, and post-filtering. The indexing stage selects only k-mers that are below a certain frequency (occurrences in the metagenomic dataset) and stores them in an inverted index called the Inexpensive K-mer Index (IKI.) This approach, inspired by the Cheap K-mer Selection step in prior work, mrFast [9], [10], filters non-informative k-mers (so-called —*expensive* k-mers) and reduces the memory footprint. The pre-filtering stage collects reads from the metagenomic dataset (i.e., the set of reads that have not been aligned to a reference) that exhibit over a predefined threshold number of inexpensive k-mers in common with the query. Then, all these candidate reads and the query are processed in the alignment stage using SSW [22], [23], a rapid SIMD implementation of the Smith-Waterman algorithm. Using five different pnealty score sets, PARMIK attempts to find the optimal alignment between the query and each candidate read. In the post-filtering step, alignment results are evaluated against user-customizable criteria, such as percentage identity and alignment size, with a focus on maximizing alignment size that meets these criteria, providing a flexible and efficient solution for metagenomic research.

Given the need to search in a large database of metagenomic sequences, there are several existing search tools. BWA (Burrows-Wheeler Aligner) [6] is commonly used for searching against a reference genome and has also been applied to metagenomic datasets [29]. BWA provides the best alignment per query. BLAST (Basic Local Alignment Search Tool) [7], [8], is a well-known multi-aligner that provides multiple alignments per query. We compared PARMIK to BLAST and BWA to find alignments between coronavirus-related samples (queries) and bat feces metagenomic datasets (reads). At a percentage identity of 90% for the alignments between queries and reads, PARMIK exceeded BLAST by achieving up to 21% higher recall. Since BLAST removes low complexity regions before indexing, BLAST’s index size was 3× smaller than PARMIK. Similarly, PARMIK removes low complexity and non-informative segments of sequences by filtering out highly frequent k-mers. After filtering highly frequent k-mers, the size of PARMIK’s index decreased by a factor of two, resulting in an index that was 1.5× larger than BLAST’s index. Moreover, PARMIK identified longer alignments faster than BLAST (up to 1.57×) when parallelizing across 32 cores. At this rate, we expect PARMIK to scale to the thousands of metagenomic environmental samples the community will generate in the coming years, allowing public health officials and researchers to fully exploit this critical resource’s capabilities to improve the analysis of metagenomic sampled data.

## II. Methods

Suppose we have a query sequence for which we want to find all the similar reads, including exact and non-exact matches, in a large metagenomic dataset with hundreds of millions of reads. To do so, we take a metagenomic dataset and create a storage-efficient inverted index (figure 3-A.) Using the index and the input queries, we find all the candidate reads that have a region of the exact match with the query (figure 3-B.) Then, each query (Q) and its candidate reads ([R]) need to go through the alignment step to find the best alignment for each (Q, R) pair (figure 3-C.) Finally, the post-filtering step should check the alignment against the user-defined criteria and improve the alignment size if possible (figure 3-D.) Detailed explanations of these steps are provided in the following sections.

**Fig. 3.**
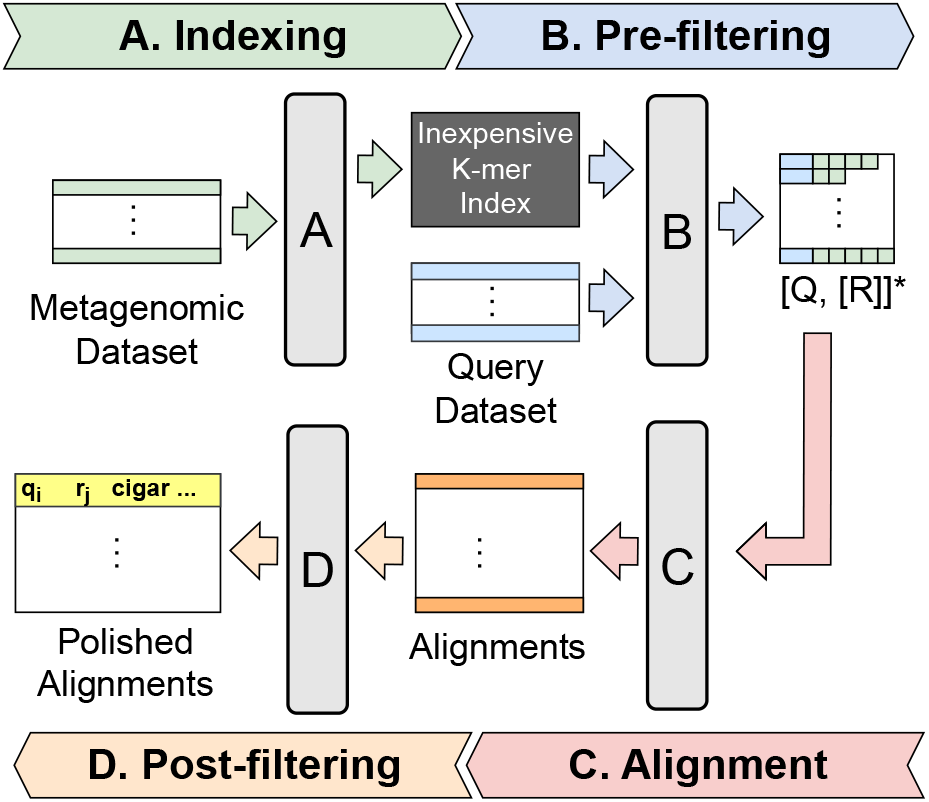
PARMIK consists of 4 steps: A. Intexing, B. Pre-filtering, C. Alignment, D. Post-filtering (*The output of the Pre-filtering step is a list of queries, each accompanied by a list of read candidates that shares exact match regions with the query.)

### A. Indexing

Metagenomic data often contains numerous sequencing errors and repetitive sequences, such as long runs of a single nucleotide, which tend to be non-informative and inflate dataset sizes significantly. Without having the reference genome, we are unable to correct errors in the metagenomic dataset, including sequencing errors. Many previous alignment methods, e.g., BLAST [7], [8], incorporate a pre-processing step to eliminate the sequencing error-related and low-complexity segments from sequences. This refinement enhances their accuracy and speed, resulting in smaller indexes.

In our study, which focuses on metagenomic datasets lacking reference genomes, we need to eliminate sequencing error-related and non-informative segments of metagenomic reads from our index. To detect these non-informative segments, we divide each read from the metagenomic dataset into k-mers and count the frequency (occurrence rate) of each k-mer. Figure 4 shows the cumulative frequency of three k-mer sizes (i.e., 9, 10, and 11) for different k-mer frequency thresholds. As the k-mer frequency threshold increases, there will be more distinct k-mers in the index. For instance, with k set to 11 and a k-mer frequency threshold of 5000, nearly 99.4% of distinct k-mers are accommodated within the index. However, with a reduction in k-mer size, keeping over 99% of k-mers in the index demands larger thresholds—specifically, 15,000 and 100,000 for k values of 10 and 9, respectively. Using this strategy, we effectively filter out non-informative k-mers that offer minimal alignment benefits, thereby preventing unnecessary inflation of the index size. Further discussion of this approach and its impact on the number of true positives, execution time, and index size is provided in section III.

**Fig. 4.**
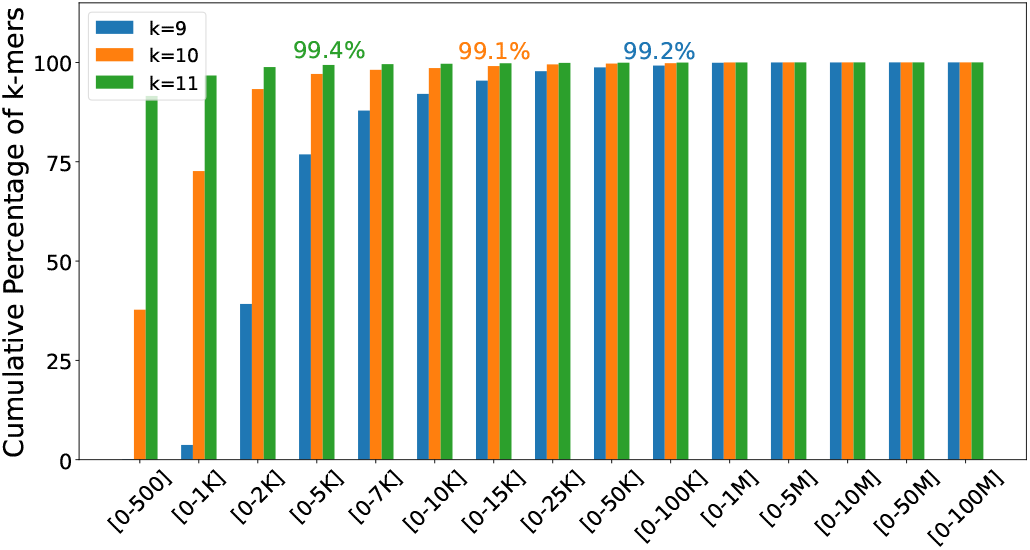
K-mer Frequency Histogram for K = [9, 10, 11].

As illustrated in Figure 5 (Indexing), all k-mers (k-1 overlapping) are extracted from the reads within the metagenomic dataset and organized into an inverted index [19]. If the k-mer frequency is higher than a pre-defined threshold (Inexpensive K-mer Threshold, IKT), we call it an *expensive k-mer*, and we do not add it to the inverted index. The inexpensive k-mers, along with their corresponding read IDs, are then incorporated into an inverted index structure called the inexpensive k-mer index (IKI). This approach is inspired by the Cheap K-mer Selection step from prior work, mrFast [9], [10]. We use the inexpensive k-mer index for the pre-filtering step.

**Fig. 5.**
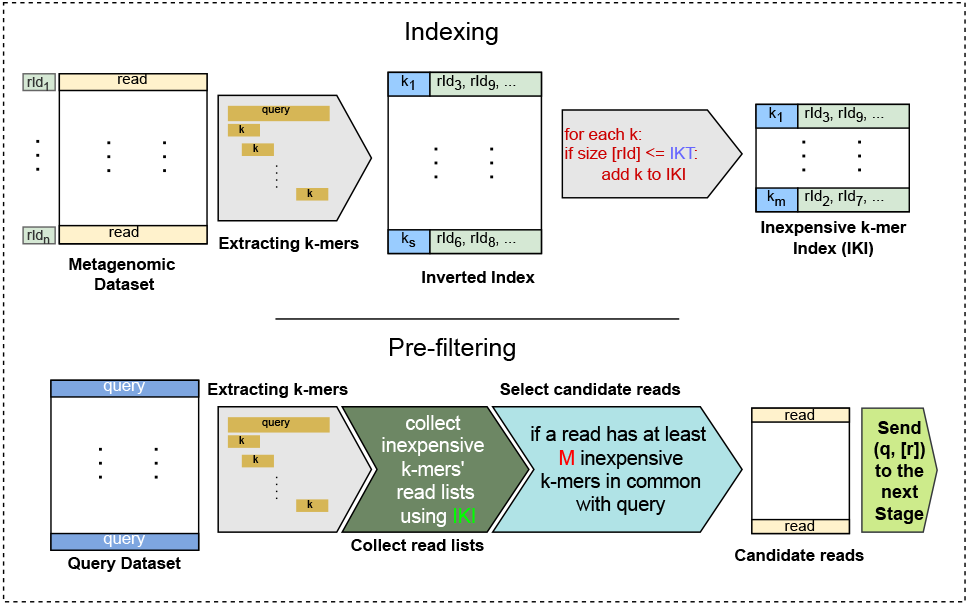
Indexing (top) and Pre-filtering (bottom) steps.

### B. Pre-filtering

PARMIK first finds reads with regions of exact matches to the query. Using k-mer matching, a well-known method [20], PARMIK ensures that two sequences share a sufficiently large common region by verifying that the number of shared k-mers surpasses a predefined threshold (M).

PARMIK uses two user-provided criteria (explained in II-D) and the k-mer size to determine the M. For example, as shown in figure 6(a), if the percentage identity is 95% and minimum alignment size is 48, the maximum number of mismatches the user expects to see in the resulting alignment is 2. If the k-mer size is 11, M is (46/11) + (46%11) = 4 + 2 = 6. To efficiently find reads similar to a query, as shown in Figure 5 (pre-filtering), we extract (k-1) overlapping k-mers from the query (and its reverse strand) and leverage IKI to identify reads sharing at least M k-mers with the query. The M k-mers do not need to be consecutive, as illustrated in Figure 6(b). Finally, we forward the query and the collected reads to the alignment step.

**Fig. 6.**
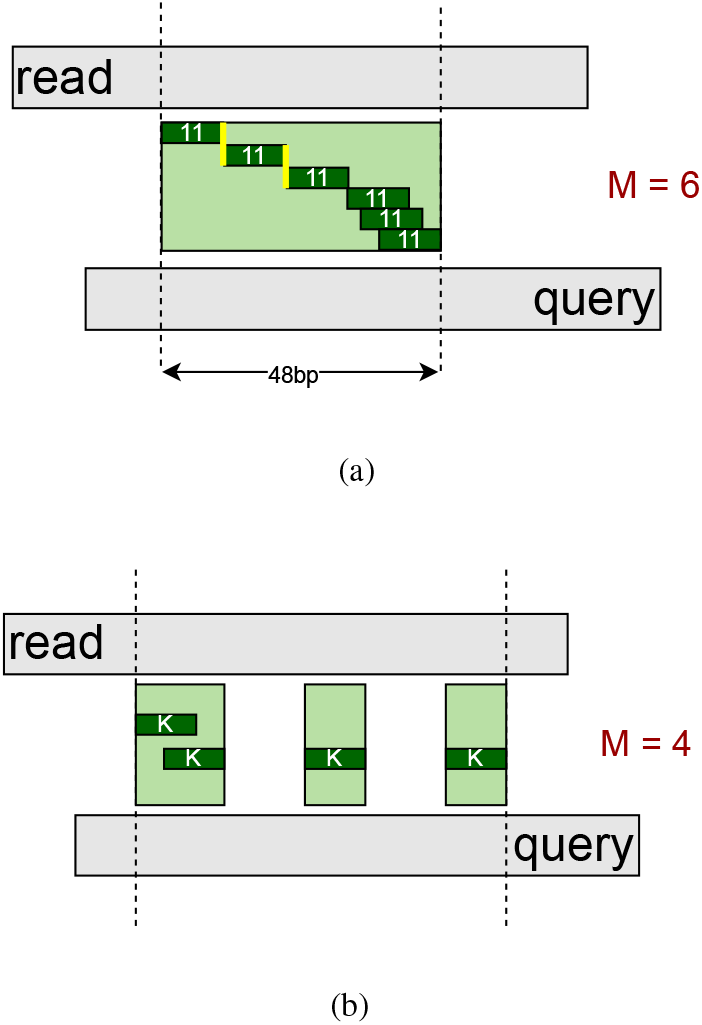
(a) An example for minimum number of k-mers shared between the read and the query sequence (M). (b) The M k-mers do not need to be consecutive.

Given the large size of data structures used for the inverted index and read ID lists in this step, achieving rapid and efficient data insertion and retrieval is imperative. We opted for the Robin map [31], implemented in C++, over the standard STL map due to its superior speed for insertions and lookups, as shown in [32].

### C. Alignment

The alignment step takes the queries and their corresponding candidate read ID lists collected during the pre-filtering stage as inputs. While Smith-Waterman [21] is a widely used method for sequence alignment, its computational cost can be prohibitive for large datasets. PARMIK uses a highly parallelized implementation of the Smith-Waterman algorithm, known as SSW. SSW incorporates an efficient SIMD (Single-Instruction-Multiple-Data) implementation of the Striped Smith-Waterman algorithm, as detailed in [22], [23].

SSW takes two sequences and the penalty scores, including *match, substitution, gap open, and gap extend*, to run the alignment. We used five distinct sets of penalty scores, as shown in the table I to check the alignments based on prioritizing different mismatch types. For instance, the penalty score set #3 prioritizes substitution (score=4) over the gap open (score=8). Then, we select the longest alignment. After identifying the optimal alignment between each read and query, the outcome is forwarded to the post-filtering step for verification against our predefined criteria.

**TABLE I.**
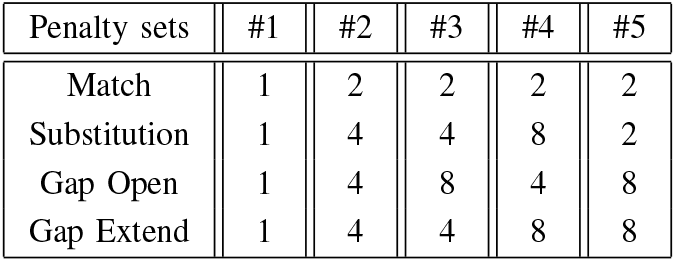
Penalty Scores.

### D. Post-filtering

Two sequences (query, read) collected from the genome sequencing may not align well, even if they are from the same genome. As shown in figure 2, the alignment region (partial match) between the query and read may be small, with regions of exact matches and some mismatches, including substitutions and indels. In this project, our goal is to identify partial matches between a read and a query. To do so, PARMIK focuses on two alignment metrics: **Percentage Identity (PI)**, which refers to the minimum percentage of the alignment containing exact matches, and **Minimum alignment Size (R)**. The results generated by PARMIK must adhere to these user-defined parameters, ensuring the percentage identity and alignment size are greater than or equal to PI and R, respectively.

Many of the alignments found by the brute force Smith-Waterman algorithm were very short (median = 11.) In our case, to help finding the pathogens for a newly discovered infection, the matches should be long enough to help us to figure out the origins of a sampled genome. Figure 7 shows as the k-mer size (size of the exact match) becomes larger, there are fewer number of reads that contain them leading to more specific read candidates for a query. Hence, our heuristics eliminate the short alignments by using the R criterion.

**Fig. 7.**
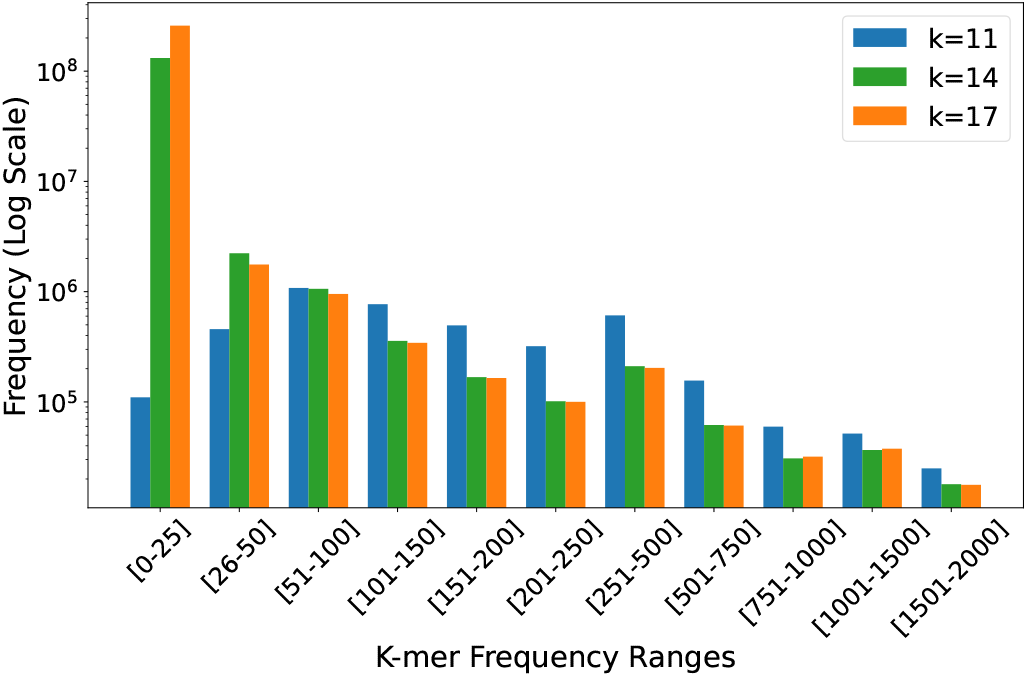
K-mer frequency variation for different k-mer sizes for Bar Feces metagenomic dataset.

If the percentage identity falls below the predefined PI, PARMIK trims the alignment from the end with fewer exact matches and recalculates the percentage identity. This process, called *Trimming*, iterates until either the percentage identity equals or surpasses PI or the alignment size decreases below the specified R threshold. After trimming, PARMIK expands the alignment by adding back the trimmed substrings, alternating sides to explore the potential for a larger alignment while ensuring the percentage identity remains equal to or greater than PI, a step we refer to as *Second Chance*. We call the trimming and second chance process as *Polishing*.

Figure 8 illustrates post-filtering for a query (Q) and read (R) where the percentage identity is initially 86% and PI=90%. To ensure alignment meets a percentage identity (*pi*) exceeding 90% criteria, PARMIK starts by trimming the read and query (❶). It counts the matches preceding the first mismatch (5) against those succeeding the last mismatch (6). Since there are fewer matches before the first mismatch, in step ❷, PARMIK trims the read and query from the front end, removing the first five exact matches followed by one mismatch. Then, PARMIK recalculates *pi*, which is (87%) still below 90%. Since the number of matches in the back (i.e., 6) end is fewer than in the front end (i.e., 12), in step ❸, six base pairs and three mismatches are trimmed from the back end. While *pi* > 90%, in step ❹, PARMIK gives a second chance to the substrings previously trimmed from the opposite end of the last trim (front end in step ❷) to potentially be added again. It recalculates *pi* to assess whether a larger alignment can be achieved while maintaining *pi* > 90%. At the end of the step, ❹, with the *pi* still surpassing 90%, PARMIK concludes this iteration by reporting the alignment as a match for the query.

**Fig. 8.**
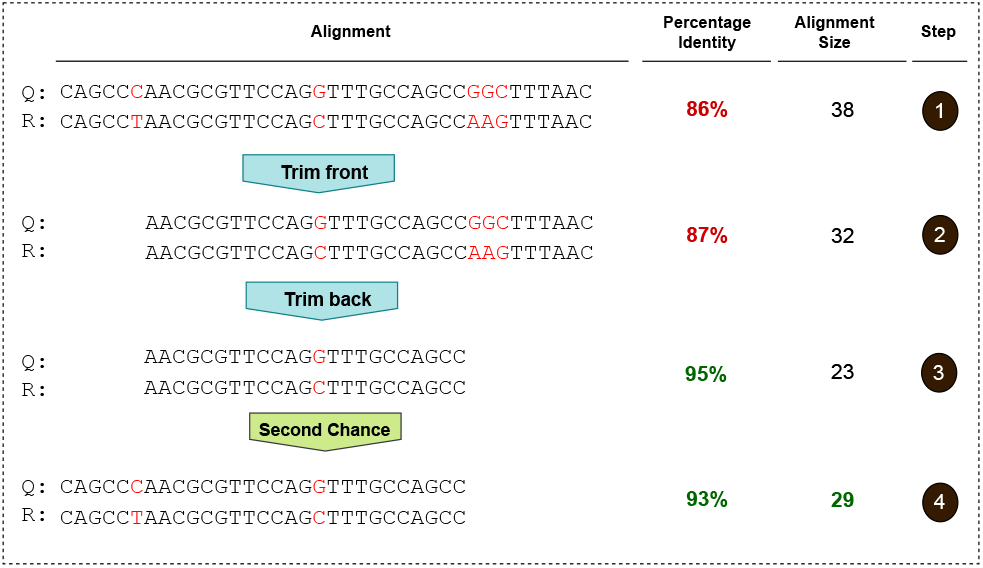
Post-filtering (PI=90%) step includes two sub-steps: (1) Trimming and (2) Second chance. Together, these sub-steps are referred to as “Polishing.”

For each query, PARMIK picks the longest polished alignment from the ten alignments generated against a read: five from the forward strand and five from the reverse strand (five penalty score sets.)

## III. Results

To evaluate PARMIK against a multi-aligner, we compared it to BLASTN, the nucleotide-focused version of BLAST+ (version 2.15.0). We set the maximum number of matches per query (max targe seqs) to an extremely high value (50,000,000) to ensure all potential matches fitting the BLAST criteria were captured. Regarding the best match per query, we compared to BWA-MEM.

### Baseline

We defined the baseline that runs the Smith-Waterman algorithm for each query against all the reads from the metagenomic dataset. If the percentage identity of the alignment result is higher than the predefined threshold (PI) and the alignment size is longer than the minimum alignment size (R), it reports the alignment for the query. For aligning each (query, read) pair and its reverse strand, we use all the penalty scores in table I and pick the alignment with the longest size.

### Input Datasets

To evaluate PARMIK, we compared the alignments from BLAST and PARMIK to the baseline for two query files against four metagenomic files (Table II.) The two query files are Sars-1 and Sars-2 (Wuhan-Hu-1.) All metagenomic datasets are collected from rainforest except SRR14381418 which is from limestone forest. The four metagenomic datasets originated from samples taken from Bat feces. We selected these metagenomic datasets based on their predicted viral load. Three of the metagenomic datasets are classified under Sarbecovirus, while SRR14381422 falls into a distinct classification of Decacovirus. We sampled 150 bp queries from assembled genomes with 149 bp consecutive overlaps. For metagenomic 150 bp reads, we removed duplicate reads and any reads containing ‘N’.

**TABLE II.** Datasets Used in Experiments.

| Dataset | Name | Size | Classification |
| --- | --- | --- | --- |
| Query | Wuhan-Hu-1 | 29 Kb | Sars-2 |
|  | SarsGenome | 29 Kb | Sars-1 |
| Metagenomic | SRR12432009 | 3.56 Gb | Sarbecovirus |
|  | SRR14381418 | 3.03 Gb | Sarbecovirus |
|  | SRR14381421 | 2.70 Gb | Sarbecovirus |
|  | SRR14381422 | 3.13 Gb | Decacovirus |

#### A. Recall & Precision

We assessed the recall and precision for PARMIK and BLAST with respect to our baseline. We ran the three tools (BLAST, PARMIK, and baseline) for the Wuhan-Hu-1 against SRR12432009 for a k-mer size of 11, a percentage identity of 90%, and Inexpensive K-mer Index contains all k-mers (IKT=all). We discuss more about the impact of different IKT later in this section.

##### 1. Recall

Figure 9(A) illustrates that throughout different minimum alignment sizes (R), the recall rate for PARMIK was higher than BLAST. This is because PARMIK found the same number or more true positives compared to the baseline, whereas BLAST lags. In addition, as the alignment size surpassed 25, PARMIK’s FN diminished compared to BLAST. Figure 9(B) focuses on the recall of the two tools for R between [21-35]. For *R ≥* 21, PARMIK exceeds BLAST by a higher recall rate, peaking at 21% for R=21. This superiority is attributed to PARMIK’s post-filtering mechanism, particularly the second chance step, which extends alignments as much as possible to maximize alignment coverage. Indeed, BLAST’s speed advantage, achieved through its heuristic approximation of the Smith-Waterman algorithm [7], is remarkable—it’s 50× faster than Smith-Waterman. However, this speed comes at the cost of precision, as BLAST tends to be less accurate than Smith-Waterman [37].

**Fig. 9.**
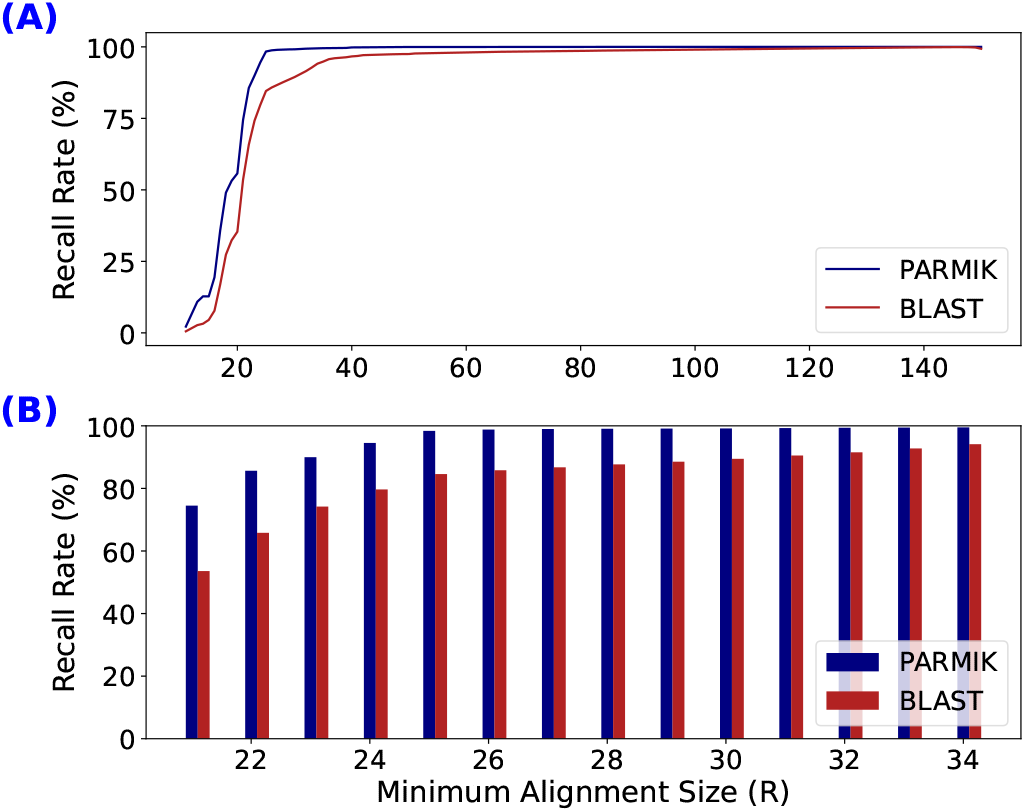
(A) Comparison of recall rates between PARMIK [K=11, M=4, PI=90%, IKT=all, polishing] and BLAST across various minimum alignment sizes (B) A zoomed-in view of the recall rate of PARMIK vs. BLAST for the range of minimum alignment sizes between 21 and 35 (Wuhan-Hu-1 vs. SRR12432009).

##### 2. Precision

Neither BLAST nor PARMIK had false positives in the experiments. Hence the precision for both of them was 1.

#### B. Speed vs. Recall

Figure 10 shows the trade-off between recall rate and execution time (minutes) for PARMIK with Inexpensive K-mer Index containing all k-mers (IKT=all), K=11, for M=4 (blue) and M=7 (green) against BLAST (red), across varying percentage identities of 85%, 90%, and 95%. Based on figure 9(B), at minimum alignment size R=30, both PARMIK and BLAST converged to high recall rates. Therefore, alignments smaller than 30 were filtered out for this experiment. While PARMIK [M=4] demonstrates a superior recall rate across various PI values, it falls short in speed compared to both BLAST and PARMIK [M=7]. As the percentage identity increases, PARMIK demonstrates reduced variability in recall rate, whereas BLAST shows increased variability in recall rate. Throughout different percentage identities, BLAST maintains a consistent runtime but shows a reduction in recall rate in some experiments. As the percentage identity increases, PARMIK’s recall rate improves, though the speed decreases. At PI = 95%, PARMIK achieves identical recall rates for M=4 and M=7, suggesting that choosing M=7 offers a good balance between recall and speed. PARMIK [M=7] is 3-5× faster than M=4 at PI=85, while the recall rate is less than 2× lower. Finally, PARMIK [M=7] has a higher recall rate and performs faster in most experiments than BLAST.

**Fig. 10.**
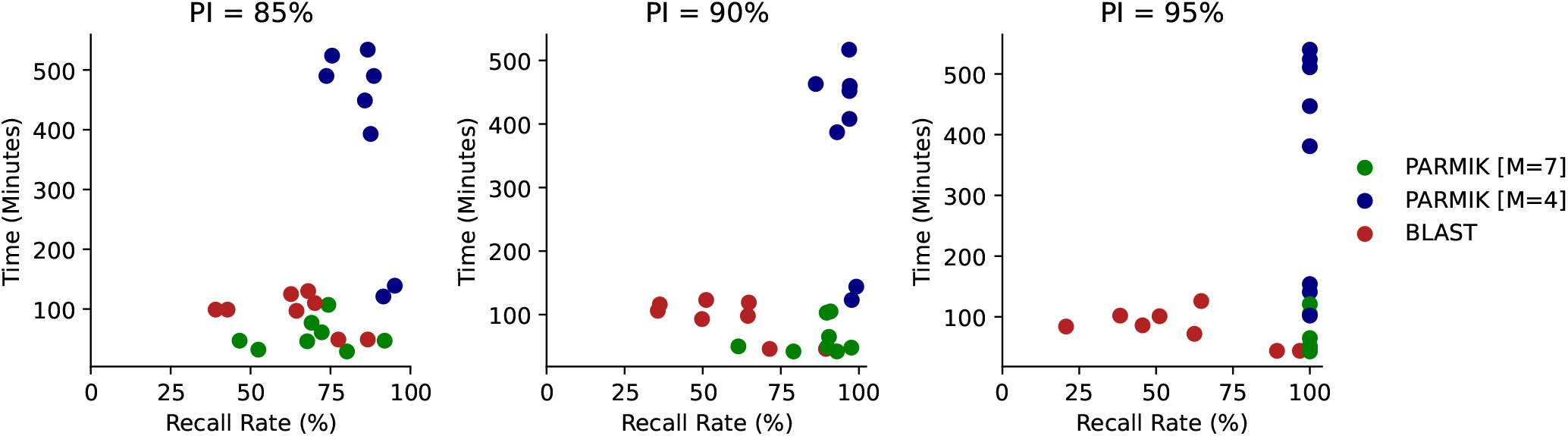
Comparison of PARMIK [K=11, IKT=all, M=7, polishing], PARMIK [K=11, IKT=all, M=4, polishing] and BLAST in terms of execution time (minutes) and recall rate (%) across all datasets for minimum alignment size of 30 and varying percentage identities of 85%, 90%, and 95%.

##### Multi-core execution

We parallelized PARMIK using OpenMP [35] and compared its performance on 32 physical cores to the parallelized version of BLAST. For Wuhan-Hu-1 against SRR12432009, under the conditions of K=11, M=4, IKT = all, and PI=90%, PARMIK and BLAST produced 30,409 and 19,227 matches per second, respectively. PARMIK was 1.57× faster than BLAST as the number of cores increased from 1 to 32.

#### C. Memory Footprint

For the metagenomic dataset SRR12432009, with K=11, M=4, and IKT = all (meaning IKI contains all k-mers, including expensive ones,) BLAST’s index is over 3× smaller than PARMIK’s index (table III.) However, we can reduce PARMIK’s memory footprint for over 2× by removing expensive k-mers from the index (more details in section III-F.)

**TABLE III.** Memory Footprint.

| Tool | Index Size (GB) |
| --- | --- |
| BLAST | 2.3 |
| PARMIK | 7.63 |

#### D. Best Match per Query

PARMIK finds all matches, while many use cases are only interested in the best match. For all three tools, including PARMIK, BLAST, and BWA, we select the alignment with the longest length among all matches to identify the best match for each query. In the experiments of aligning Wuhan-Hu-1 against SRR12432009 with a k-mer size of 11, IKT = all, and a percentage identity threshold of 90%, PARMIK successfully found at least one match for all queries. Figure 11(a)(green bars) demonstrates that for the queries that BLAST failed to identify any alignments, 507 (1.7%) of the total queries, PARMIK found alignments ranging between 20 to 60 bp. Similarly, figure 11(b)(green bars) shows that PARMIK identified alignments of size up to 150 bp for the cases that BWA found no alignments for the query, 3076 (10.3%) of the total queries.

**Fig. 11.**
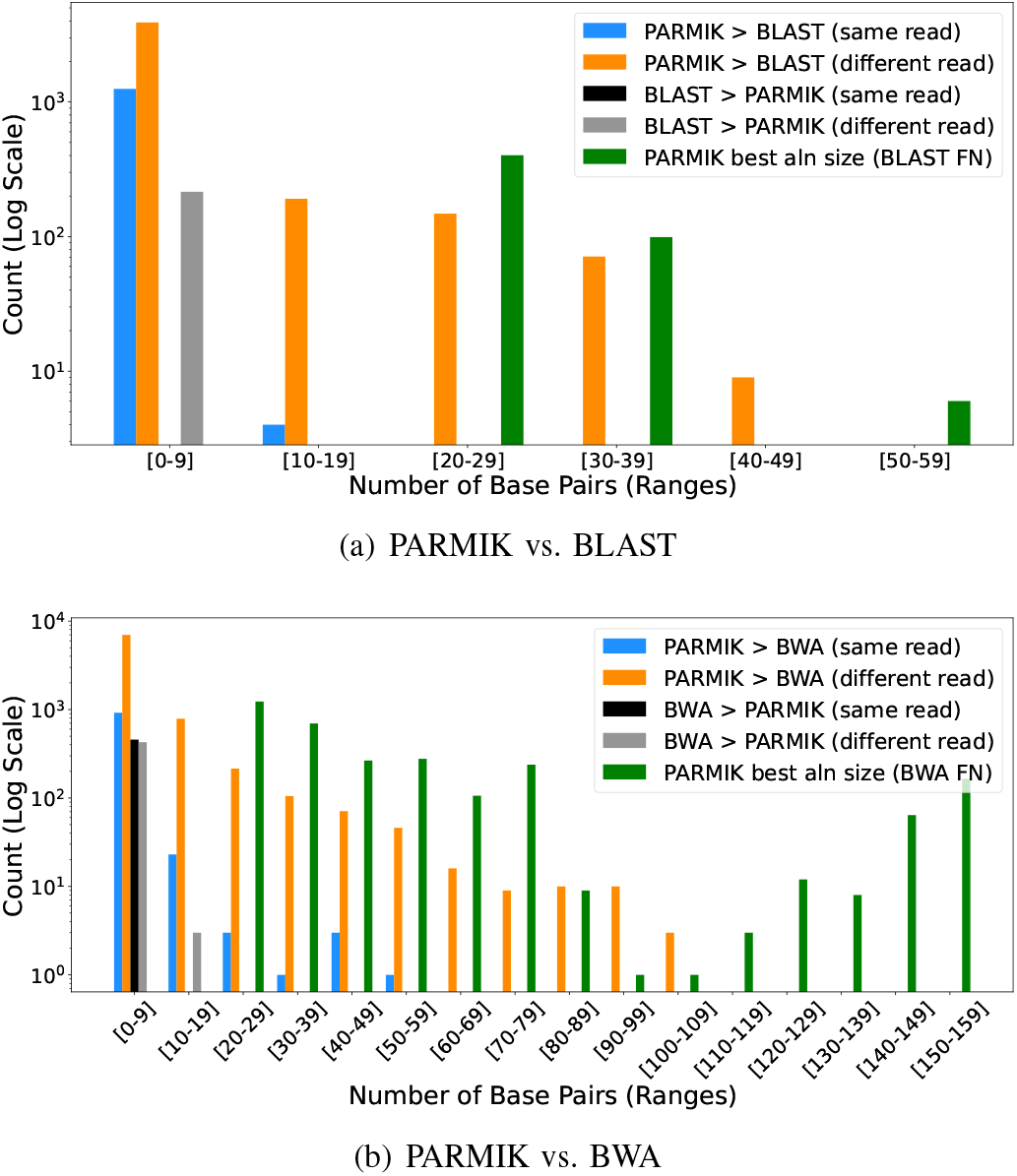
Histogram of alignment length difference for the best match per query (PARMIK vs. BLAST/BWA.) “Different read” or “same read” means PARMIK and BLAST/BWA found different or the same best read per query (Wuhan-Hu-1 vs. SRR12432009).

As shown in figure 12, BLAST and BWA identified at least an alignment for 29,247 and 26,678 out of 29,754 queries, respectively. When comparing the best match for each query between PARMIK and BLAST as well as BWA, for 99.2% and 96.6% of queries, PARMIK identifies alignments of longer or the same alignment size per query compared to BLAST and BWA, respectively. However, there is a small number of queries, 0.8% and 3%, that BLAST and BWA outperform PARMIK in terms of alignment size of best match. We checked alignments where BLAST and BWA outperformed PARMIK. For BLAST, all such cases involved short alignment sizes, typically around 20 bp. For BWA, these alignments exhibited a median of 1 bp fewer soft clips compared to PARMIK. The slightly higher number of soft clips in PARMIK results from the SSW algorithm, which extends the alignment until it reaches a score of 0 at either end, then adds soft clips for the 0-scored ends of the alignment.

**Fig. 12.**
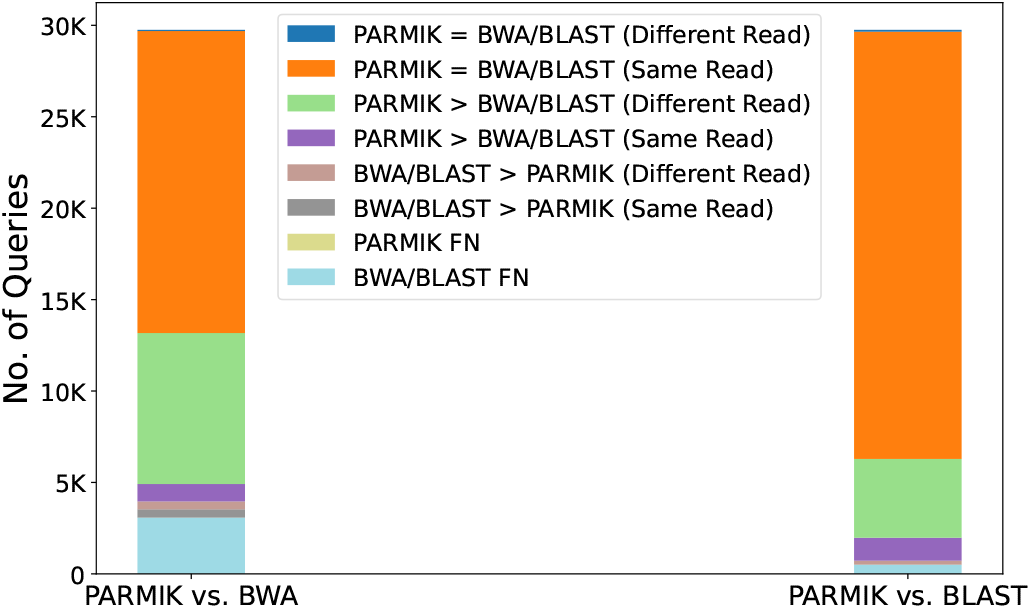
Comparing best match per query (PARMIK vs. BLAST/BWA). Different (Same) read means PARMIK and BLAST/BWA found different (similar) read for the query (Wuhan-Hu-1 vs. SRR12432009).

Among the cases where PARMIK finds a similar alignment size to BLAST and BWA, it identifies the same read as BLAST for 78.5% of the times and the same read as BWA for 55.5% of the times. Additionally, PARMIK outperforms BLAST and BWA by identifying a longer alignment for 18.7% and 31% of queries, respectively. Figures 11(a) and 11(b) show the number of cases that each tool found better alignment than the other tool, measured by the difference in the length of alignment found by the two tools. PARMIK outperforms BLAST and BWA by identifying longer alignments with a median of 2 and 3 more bp, respectively.

#### E. Impact of Polishing in the Post-filtering Step

To assess the polishing technique’s efficacy detailed in Section II-D, we employed two variants of the PARMIK tool: one equipped with polishing in the post-filtering step and another without this capability. For this analysis, we used the Wuhan-Hu-1 dataset against SRR12432009 using a k-mer size of 11 and a 90% percentage identity threshold.

Figure 13(a) depicts the relationship between the preliminary alignment size (*x-axis*) and the alignment size after polishing (*y-axis*). Notably, every data point either maintained the same size as the preliminary alignment or exceeds it after polishing since there are no points below the line *y* = *x*. Furthermore, in some cases, the alignment size significantly increased after polishing.

**Fig. 13.**
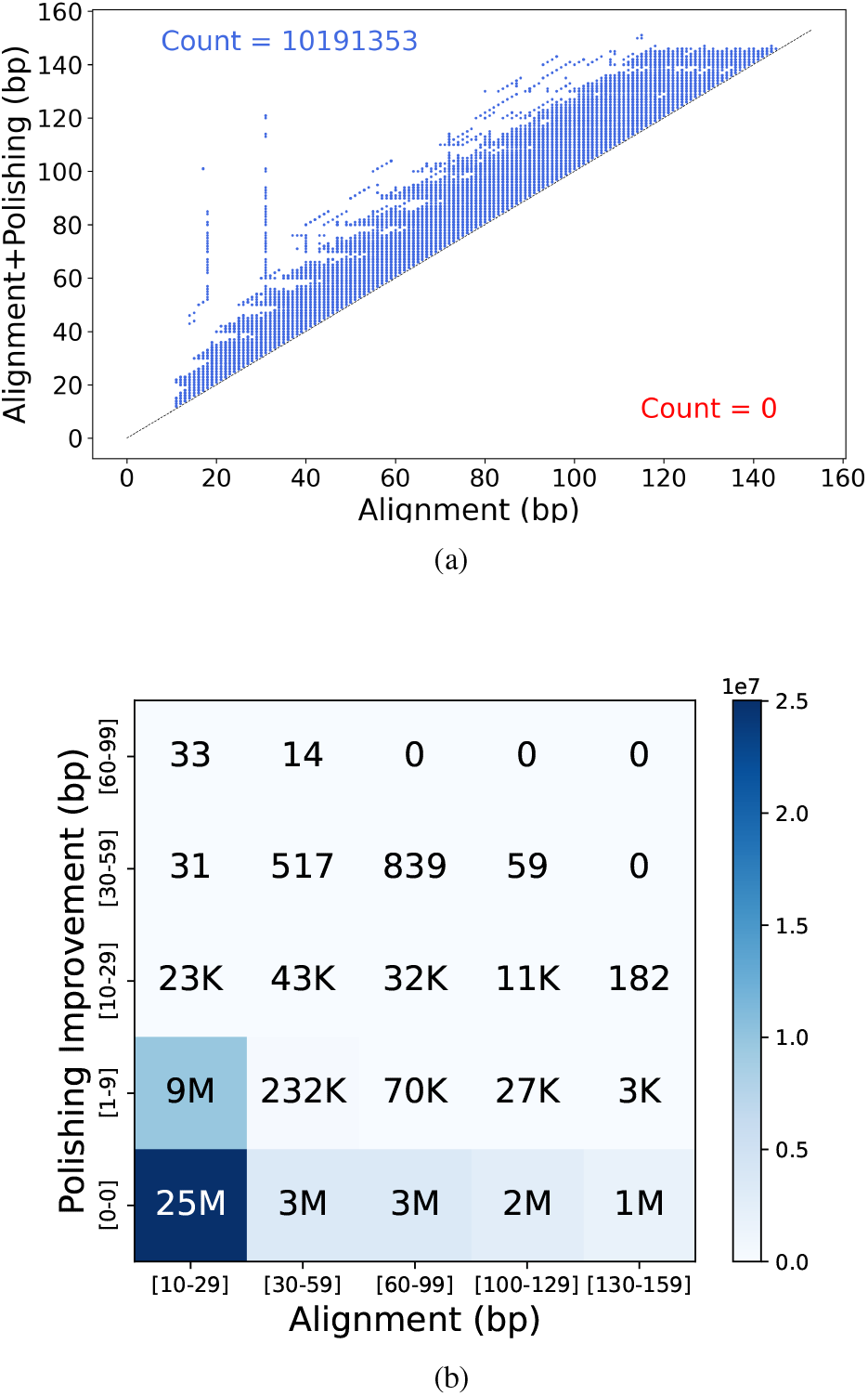
Impact of using Polishing on PARMIK’s Alignment Size. (a) Alignment size improvement due to polishing per alignment. The x-axis shows the preliminary alignment size, and the y-axis shows the alignment size after polishing. (b) Heatmap of the histogram of polishing improvements. Each box shows the count of the preliminary alignment size range (x-axis) improved to the range shown on the y-axis. For example, 232K alignments of size between [30-59] bp improved to a range of [1-9] bp. An alignment improvement range of [0,0] signifies no improvement achieved from using the polishing technique (Wuhan-Hu-1 vs. SRR12432009).

Figure 13(b) shows the extent to which polishing enhances alignment size. The *x-axis* represents ranges of preliminary alignment sizes, while the *y-axis* depicts ranges of alignment improvement after using the polishing technique. An alignment improvement range of [0,0] signifies no improvement achieved from using the polishing technique, comprising 75% of the observed cases. This figure shows the significant impact of polishing during the post-filtering step, resulting in an increase in alignment size for over 10 million or 25% of the alignments. Moreover, polishing enhanced alignment size 3 to 9 times for shorter alignments, such as those within the range [10-29], extending up to [60-99].

Figure 14 illustrates an example that an initial alignment of size 18 improves to 81 after polishing. PARMIK uses five different penalty score sets and performs alignments on both the forward and backward strands of the read and query (10 alignments per query.) After post-filtering, PARMIK picks the longest alignment size among 10 alignments for this read and query. The two alignments in the figure are from the two different strands of the query, which indicates that the strand identified with the polishing technique enabled did not pass the post-filtering step when the polishing technique was disabled. In other words, a preliminary alignment that is discarded for a particular penalty score set may be accepted and significantly improved in size (e.g., from 18 to 81) when PARMIK’s polishing technique is enabled.

**Fig. 14.**
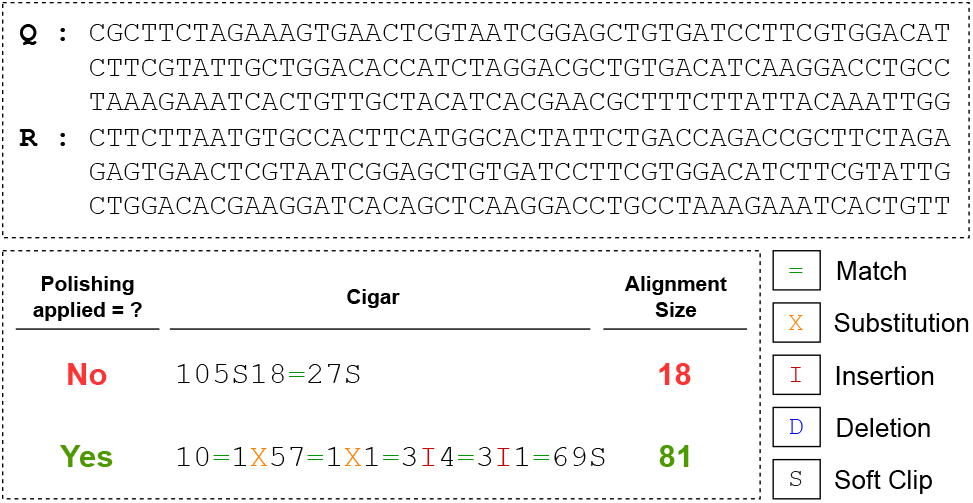
An example of improved alignment size due to polishing.

#### F. Impact of using Inexpensive K-mers

Metagenomic data often contain sequencing errors and non-informative, highly repetitive sequences, which leads to larger datasets and needs more memory. Pre-processing steps in previous alignment methods, such as BLAST, address this by filtering out these errors and low-complexity segments, reducing the memory footprint. To address this, we filter expensive k-mers resulting in smaller memory footprint.

Figure 4 shows that for K=11 and IKT=5000, 99% of k-mers were included within the IKI. Figure 15-A illustrates the impact of excluding expensive k-mers with *IKT* > 5000 on recall rate. For all minimum alignment sizes, the recall rate of PARMIK [IKT=5000] exceeded that of BLAST but fell short compared to PARMIK [IKT=all], particularly for 25 < *R* < 35 (Figure 15-B). When *R* > 35, the recall rate of PARMIK with a 5000-threshold became close to the PARMIK [IKT=all]. Additionally, the size of the index decreased by more than 2×, from 7.63 GB to 3.47 GB, and the execution time was reduced by 12% when using the 5000-threshold. Figure 15-C shows that PARMIK [IKT=all] outperforms all others, including the Baseline, in terms of the number of true positives across various values of R. Figure 15-D presents a histogram of the alignment size differences between PARMIK [IKT=all, polishing] and the Baseline, highlighting that PARMIK identifies more longer alignments thanks to the polishing step.

**Fig. 15.**
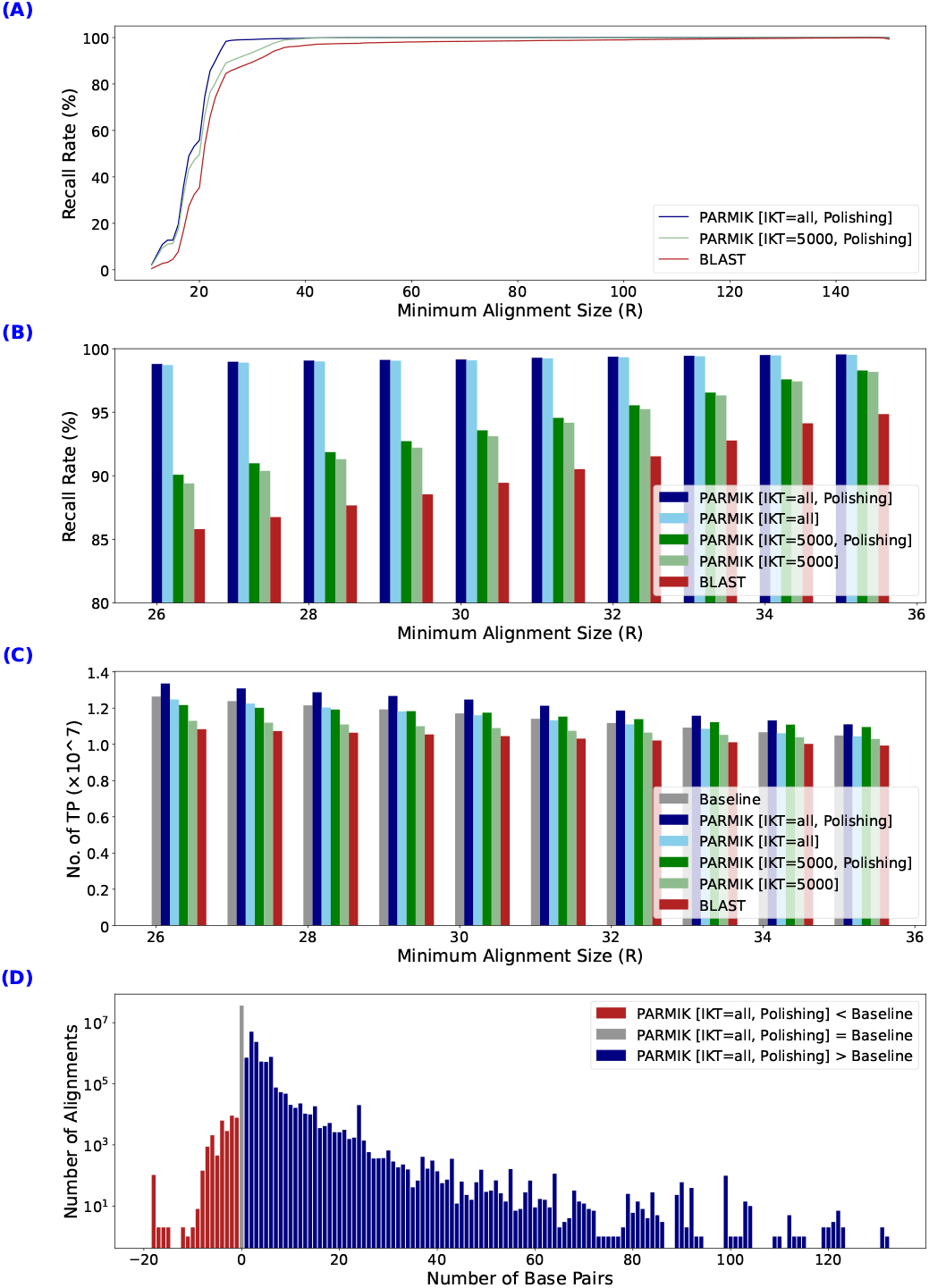
(A, B) Comparison of recall rates between PARMIK [IKT=all] (with and without polishing), PARMIK [IKT=5000] (with and without polishing), and BLAST across various minimum alignment sizes (C) Comparison of true positive rates between PARMIK [IKT=all] (with and without polishing), PARMIK [IKT=5000] (with and without polishing), and BLAST across various minimum alignment sizes (D) Comparison of alignments sizes between PARMIK PARMIK [IKT=all] and Baseline for the true positives they found (Wuhan-Hu-1 vs. SRR12432009).

**Fig. 16.**
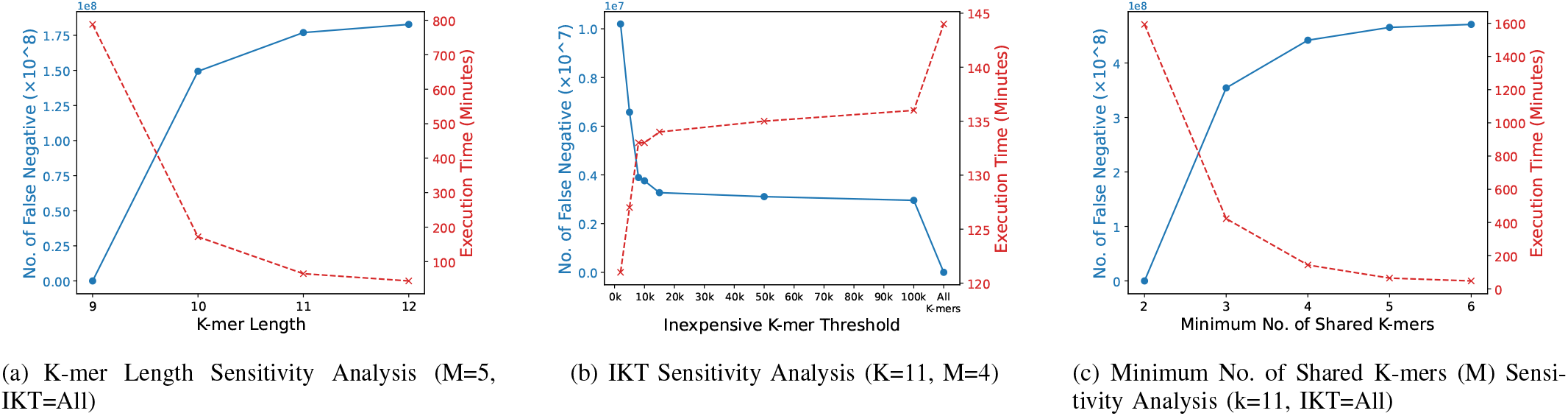
Sensitivity Analysis of PARMIK: The false negative values are normalized to the highest IKT in 16(b) (when IKI has all the k-mers) or the lowest value in 16(a) (i.e., 9) and 16(c) (i.e., 2). The median alignment size of the false negative cases is 13 for 16(a) when k=11, 15 for 16(b) when IKT=5000, and 12 for 16(c) when M=4 (Wuhan-Hu-1 vs. SRR12432009).

PARMIK containing k-mers with threshold over 5000 missed 8,437 (0.0004%) matches due to filtering out expensive k-mers. We analyzed these expensive k-mers in the reads and found that 94% of them would not be considered expensive if we increased the IKT to a higher value, i.e., 10,000. We collected the most expensive k-mers with a frequency greater than one million in the missed matches and constructed their phylogenetic tree using Clustal Omega [36]. The analysis revealed that these expensive k-mers are highly clustered, forming 8 distinct clusters. In each cluster, the k-mers are quite similar, partly or totally. Figure 17 illustrates a cluster of 25 k-mers, each with a frequency over 1 million, showing their remarkable similarity. Additionally, the complexity in 72 out of 150 of these k-mers is less than 50%. The analysis also showed that the top three most expensive k-mers are Illumina global adapters which are required for the sequencing step but not informative for the analysis step.

**Fig. 17.**
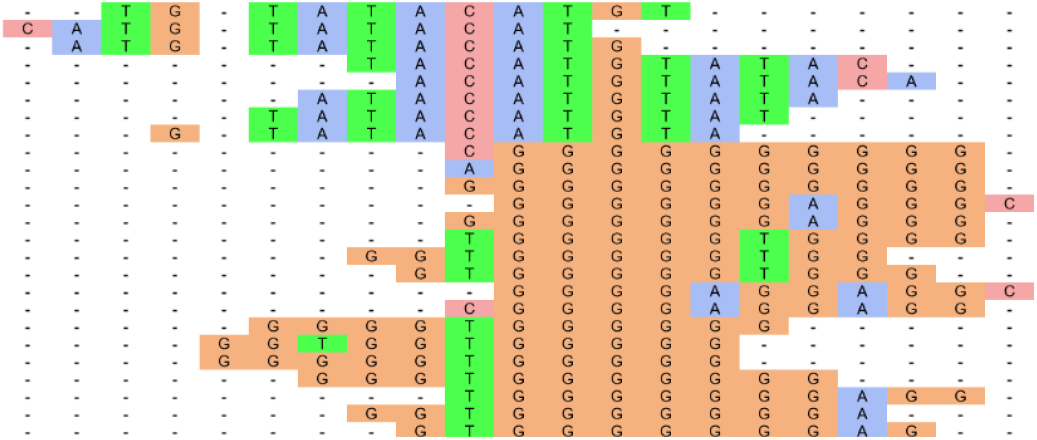
Cluster of 25 highly-frequent expensive k-mers of size 11, each exceeding 1 Million in frequency.

Eliminating expensive k-mers from the IKI not only decreases the memory footprint and execution time but also maintains the same level of TP comparable to PARMIK [IKT=all], which includes all k-mers. To show this, we compared the recall rate of the PARMIK with K=11, M=4, for IKT=all and IKT=5000, and polishing against BLAST across all different datasets. Figure 18 illustrates that PARMIK [IKT=all, polishing] achieves superior performance, followed by PARMIK [IKT=5000, polishing], both of which outperform BLAST. The impact of filtering expensive k-mers on recall rate was less than 5% throughout all datasets. The impact of only using inexpensive k-mers on IKI size and speed is anticipated to be significantly greater when comparing a query against multiple large metagenomic datasets, which is more abundant in non-informative, expensive k-mers.

**Fig. 18.**
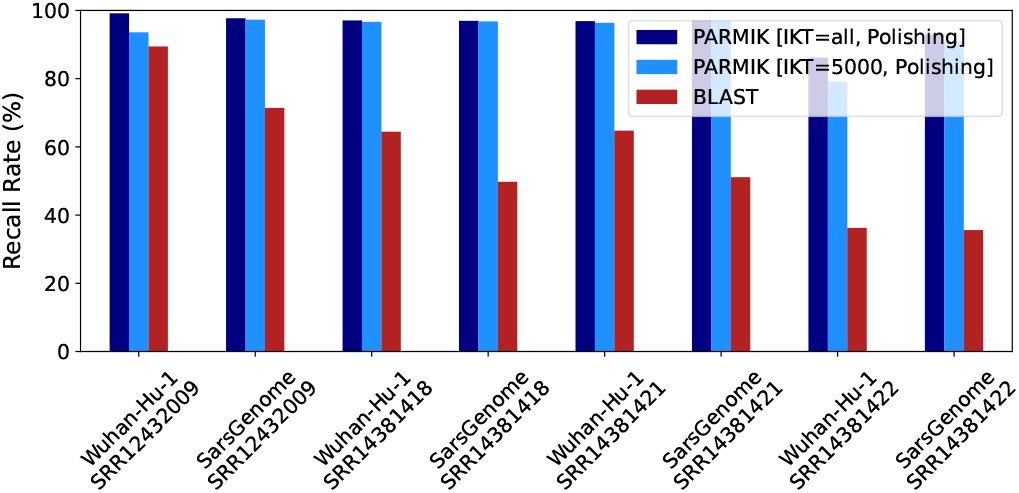
Comparison of recall rates for PARMIK [IKT=5000, polishing], PARMIK [IKT=all, polishing], and BLAST across all datasets, evaluated at R=30.

#### G. Sensitivity Analysis of PARMIK

To evaluate the sensitivity of PARMIK to its various parameters (figure 16), we changed one parameter while keeping the others fixed and measured the false negative (blue diagram) compared to the baseline and the execution time (red diagram.)

Figure 16(a) shows the sensitivity analysis of PARMIK to k-mer length. We set M=5 and IKT=0 and change the k-mer length from 9 to 12. Increasing the k-mer length led to a rise in the number of false negatives and a fall in execution time. All false negative numbers were normalized to K=9 (subtracted by its false negative.) There was a sharp increase of about 140 million in false negatives when the k-mer length was changed from 9 to 10.

We also evaluated the sensitivity analysis of PARMIK to Inexpensive K-mer Threshold (IKT) by setting K=11 and M=4. Figure 16(b) shows that (all false negative numbers were normalized to the case we have all k-mers in the IKI) as IKT increased, the number of false negatives decreased, and the execution time increased. This effect was particularly more significant when IKT changed from 2k to 10k.

We assessed the sensitivity analysis of PARMIK with respect to the minimum number of shared k-mers (M) by setting K=11 and IKT=0. Figure 16(c) shows that (all false negative numbers were normalized to M=2) as M increased, the number of false negatives increased, and the execution time decreased. This effect was particularly more significant when M changed from 2 to 3.

## IV. Related work

### A. BLAST

The Basic Local Alignment Search Tool (BLAST) [7], [8] is a widely utilized bioinformatics tool for sequence comparison and analysis. BLAST was first introduced by Altschul et al. in 1990 and has since become a cornerstone in molecular biology research. Numerous studies have focused on enhancing BLAST’s algorithmic efficiency, accuracy, and applicability to various biological data types. Although BLAST generally outpaces most Smith-Waterman implementations in speed, it doesn’t assure optimal alignments as the Smith-Waterman algorithm does. Instead, BLAST employs a heuristic method to search for high-scoring sequence alignments between the query and database sequences, approximating the Smith-Waterman algorithm [33]. BLAST+ [25] is the most recent version of BLAST, which offers improved performance and additional functionalities. We compared PARMIK with BlastN, the nucleotide alignment variant of BLAST+ (version 2.15.0), evaluating various metrics, including sensitivity and speed.

### B. BWA

Burrows-Wheeler Aligner (BWA) [6] is a prominent read alignment tool that accounts for about 30-40% time of the reference-guided assembly pipeline. At its core, BWA employs the Burrows-Wheeler Transform [18] (BWT) and the Ferragina-Manzini index [17] (FM-index) to facilitate efficient alignment processes. The BWT [18] is a reversible transformation of the reference genome that enhances compression and accelerates search operations. The FM-index, derived from the BWT, further optimizes the ability to locate specific sequences within the reference genome swiftly. BWA employs a seed-and-extend strategy where it first generates short substrings (seeds) from the reads. The algorithm then extends these seeds using dynamic programming, incorporating a variant of the Smith-Waterman [21] algorithm for accurate and sensitive alignment. The algorithm is robust to sequencing errors and applicable to a wide range of sequence lengths from 70 bp to a few megabases. For mapping 150 bp sequences, BWA-MEM (BWA’s latest variant) shows better performance than several state-of-the-art read aligners to date. Since BWA is not a multi-aligner—meaning it is designed to identify only the best (longest) alignment per query rather than multiple alignments—we compared PARMIK to BWA (BWA-MEM variant) solely in terms of identifying the best alignment.

### C. mrFast

Another notable tool in sequence alignment is mrFast [9], a storage-efficient and versatile short-read alignment tool. The tool efficiently handles large-scale sequencing data, making it particularly valuable for projects demanding accurate alignment, such as genome-wide association studies (GWAS) and clinical genomics. mrFast integrates FastHASH [10], an indexing scheme that boosts performance in seed-and-extend hash table-based read mapping algorithms. FastHASH maintains high sensitivity while introducing two key techniques: Adjacency Filtering (AF) and Cheap K-mer Selection (CKS). Our Inexpensive K-mer Collection step was inspired by the CKS method used in mrFast. We initially tried both the binary and docker versions of mrFast but found them to be prohibitively slow. As a result, we decided to exclude mrFast from our comparison.

### D. Bowtie

Bowtie [11] is a genome alignment tool known for its speed and memory efficiency. It aligns short DNA reads to large genomes using Burrows-Wheeler indexing, aligning 25 million reads per CPU hour for the human genome with only 1.3 GB of memory. Bowtie’s quality-aware backtracking algorithm allows for mismatches, improving alignment accuracy. It also supports multiple processor cores for faster alignment. The first version of Bowtie does not support gapped alignment. Bowtie2 [12], released in 2012, addresses these limitations and adds support for gapped, local, and paired-end alignment modes. Similar to BWA [6], Bowtie2 uses the FM index. Since Bowtie2 and BWA are working similarly, we chose BWA for comparison with PARMIK.

### E. Kraken

Kraken [13] stands as a well-known tool for fast and accurately assigning taxonomic labels to metagenomic DNA sequences. Unlike its predecessors, which were slower and computationally intensive, Kraken excels in speed and precision. Through exact k-mer alignment, it achieves classification accuracy comparable to the fastest BLAST programs. In its fastest mode, Kraken can classify 100 base pair reads at a rate of over 4.1 million reads per minute, outpacing alternative tools by a large margin. We opted not to compare PARMIK with Kraken because Kraken is a taxonomy classification tool that does not provide alignment coordinates.

### F. Centrifuge

Centrifuge [15] is a microbial classification engine for desktop computers, offering rapid and precise labeling of genetic reads. Its indexing scheme, based on BWT [18] and FM-index [17], allows for accurate species quantification with a small index size (4.2GB for 4078 bacterial and 200 archaeal genomes). The FM-index empowers Centrifuge to swiftly search k-mers of varying lengths, achieving speeds akin to traditional k-mer table indexing algorithms. Centrifuge initiates with a brief exact match (minimum 16 base pairs) and extends the match to its maximum extent. Moreover, Centrifuge’s space-optimized indexing can encompass the extensive NCBI nonredundant nucleotide sequence database (109 billion bases) with only a 69 GB index size, a notable improvement over k-mer-based approaches. We did not compare PARMIK with Centrifuge since it does not give the alignment coordinates.

### G. Themisto

Pseudoalignment is a fast and cost-effective method for determining if a read matches a reference sequence without providing exact alignment coordinates. Themisto [16], an example of a pseudoalignment method, is a scalable colored k-mer index tailored for extensive collections of microbial reference genomes, accommodating both short and long-read data efficiently. It indexes 179,000 Salmonella enterica genomes within 9 hours, resulting in a 142 GB index. In contrast, leading competitors such as Metagraph [38] and Bifrost [39] only managed to index 11,000 genomes within the same timeframe. Notably, in pseudoalignment, Themisto significantly outperforms these alternatives, showcasing either superior speed or memory efficiency. Additionally, Themisto achieves higher recall on Nanopore read sets, emphasizing its superior pseudoalignment quality. Themisto is a pseudoalignment method that does not provide exact alignment coordinates, making it unsuitable for comparison with PARMIK.

## V. Conclusions

Metagenomic sampling is important for pandemic preparedness, providing a proactive means to detect potential pathogens and implement timely interventions. However, analyzing diverse metagenomic environments, where many reads lack matches in reference databases, remains a bottleneck despite rapid DNA read generation through high-throughput sequencing technologies. Existing tools may fail in detecting emerging or undiscovered pathogens, necessitating advanced search tools for efficient and precise identification of relevant reads. We proposed PARMIK, a partial matching tool to identify a partial match between two sequences where the boundaries of the two sequences do not necessarily align, and the overlapping region can be small. Our study demonstrates that at a 90% percentage identity threshold, PARMIK surpasses BLAST achieving up to 21% higher recall than BLAST. Additionally, PARMIK outperforms BLAST in identifying longer alignments and retrieves them much faster, with up to 1.57× faster end-to-end execution time compared to BLAST when parallelizing across 32 cores. Moreover, by filtering out highly frequent (expensive) k-mers, PARMIK operates 12% faster than BLAST while reducing the index size by more than 50%. With multiple large metagenomic datasets, where expensive k-mers are more prevalent, we anticipate a greater impact of excluding expensive k-mers on the IKI size and speed. Hence, PARMIK is expected to scale to accommodate thousands of metagenomic environmental samples, empowering public health officials and researchers to fully leverage its capabilities to improve the analysis of metagenomic sampled data.

## VI. Acknowledgement

This work was supported in part by PRISM, one of six centers in JUMP, a Semiconductor Research Corporation (SRC) program, sponsored by MARCO and DARPA.

## Footnotes

* The letter “k” refers to the length of the substring.

